# Yeast [FeFe]-hydrogenase-like protein Nar1 can bind not only two [4Fe-4S] clusters but also a [2Fe-2S] cluster

**DOI:** 10.1101/2025.03.25.644927

**Authors:** Joseph J. Braymer, Lukas Knauer, Jason C. Crack, Jonathan Oltmanns, Melanie Heghmanns, Jéssica C. Soares, Nick E. Le Brun, Volker Schünemann, Müge Kasanmascheff

## Abstract

Nar1 is an essential eukaryotic protein proposed to function as an iron-sulfur (Fe/S) cluster trafficking factor in the cytosolic iron-sulfur assembly (CIA) machinery. However, such a role has remained unclear due to difficulties in purifying adequate amounts of cofactor-bound protein. The [FeFe]-hydrogenase-like protein has two conserved binding sites for [4Fe-4S] clusters, one of which is predicted to be a labile site for cluster transfer to downstream targets. Here, we report a new preparation procedure for Nar1 that facilitated studies by UV-Vis, EPR, and Mössbauer spectroscopies, along with native mass spectrometry. Nar1 recombinantly produced in *E. coli* contained a [4Fe-4S] cluster, bound presumably at site 1, along with an unexpected [2Fe-2S] cluster bound at an unknown site. Fe/S reconstitution reactions installed a second [4Fe-4S] cluster at site 2, leading to protein with three Fe/S cofactors. Strikingly, one [4Fe-4S] cluster was rapidly destroyed by molecular oxygen, potentially linking Nar1 oxygen sensitivity to phenotypes observed previously *in vivo*. These advances now allow for the pursuit of *in vitro* Fe/S cluster transfer assays, which will shed light on Fe/S trafficking by CIA components and how they may facilitate the insertion of [4Fe-4S] and potentially [2Fe-2S] clusters into target proteins in the cytosol.

---

The generation, trafficking, and insertion of [4Fe-4S] clusters into cytosolic and nuclear [4Fe-4S] proteins is a fundamental reaction required for the proper maturation of essential DNA/RNA processing enzymes, amongst many others.^[1–2]^ These functions are carried out by the cytosolic iron-sulfur (Fe/S) cluster assembly (CIA) machinery in eukaryotes, which can consist of up to 13 proteins and is also fully dependent on the mitochondrial Fe/S cluster assembly (ISC) system.^[3–6]^ In the model of [4Fe-4S] cluster biogenesis in the cytosol, tetranuclear clusters are generated on the yeast scaffolding proteins Cfd1 and Nbp35 via the assistance of reducing power from the electron transfer complex involving NADPH, Tah18, and Dre2.^[7–8]^ The generated [4Fe-4S] clusters are then trafficked to Nar1^[9–11]^, which connects further to Cia1, Cia2, and Mms19 of the cytosolic targeting complex (CTC) that is responsible for the final insertion of Fe/S clusters into target proteins.^[12–13]^

Nar1 is a structural [FeFe]-hydrogenase homolog that lacks hydrogenase activity and may be both a CIA component and a potential target of the CIA machinery, as it contains two conserved [4Fe-4S] binding sites (Figure 1) along with a C-terminal tryptophan for targeting to the CTC (Figure S1).^[9–10, 13]^ Site 1 in the C-terminal domain is homologous to the [4Fe-4S]H cluster binding site of the H-cluster in [FeFe]-hydrogenases (Figure 1). A complete H-cluster including the unique 2Fe cluster, [2Fe]H, is not found in eukaryotic yeast and human Nar1 homologs.^[10]^ Site 2 of Nar1 contains four conserved cysteine residues comparable to the FS4A cluster binding site in [FeFe]-hydrogenases (Figure 1).^[15]^ Previous attempts to purify Nar1 from several organisms have yielded protein with predominately only one [4Fe-4S] cluster in site 1, precluding a thorough biochemical and biophysical analysis of the protein.^[9, 11, 16]^ While other CIA components have been isolated and biophysically characterized ^[12, 17–20]^, the inability to produce mature Nar1 has also prevented the development of *in vitro* CIA pathway assays of the type established for the early and late ISC machineries. ^[21–22]^

**Figure 1.**
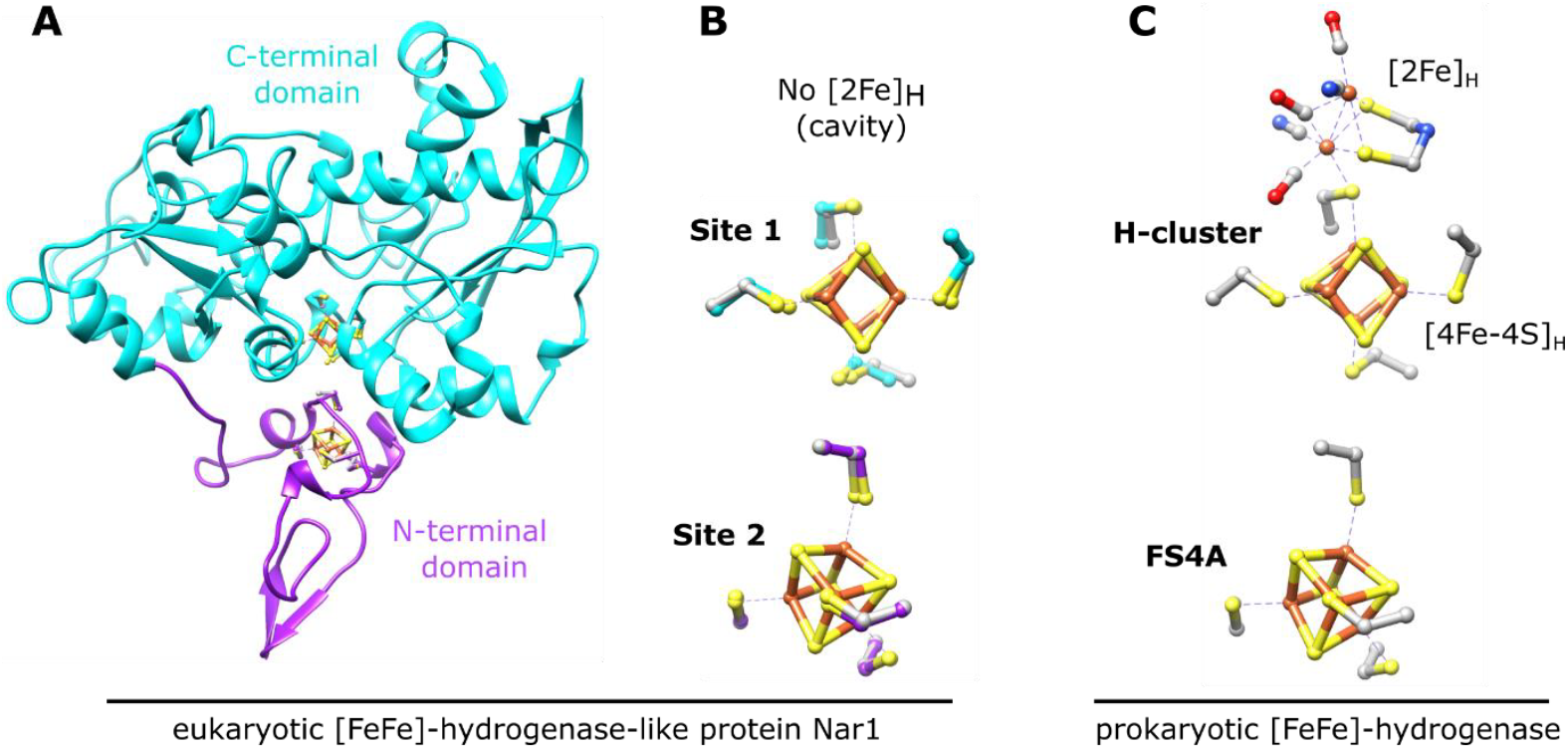
Nar1 is predicted to bind two [4Fe-4S] clusters based on homology with prokaryotic [FeFe]-hydrogenases. A) Predicted protein structure of Nar1 showing the N-terminal ferredoxin-like domain (magenta) and the C-terminal domain (cyan). The yeast Nar1 sequence was threaded through the [FeFe]-hydrogenase from *Clostridium pasteurianum* (PDB 1FEH) using SwissModel^[14]^ and the [4Fe-4S] clusters from 1FEH were overlayed onto the modelled Nar1 structure. B) Coordination environments of the two predicted [4Fe-4S] cluster binding sites of Nar1 as shown in (A). Corresponding coordinating cysteines from 1FEH are shown in grey. C) The H-cluster, composed of [2Fe]_H_ and [4Fe-4S]_H_, and the proximal [4Fe-4S] cluster (FS4A) of [FeFe]-hydrogenases are shown (PDB 4XDC).

In an attempt to produce mature Nar1, an *E. coli* host strain with a functional SUF machinery (Suf^++^)^[23]^ was utilized to express N-terminally His-tagged and C-terminally Strep-tagged Nar1 from *S. cerevisiae* (supplemental methods). Recombinant Nar1 was isolated and purified under anaerobic conditions as a predominately monomeric, brown-colored protein solution with UV-vis absorption features at 320 and 420 nm indicative of [2Fe-2S] and/or [4Fe-4S] clusters (Figure S2).^[24]^ Native mass spectrometry confirmed a monomeric, folded state of Nar1 (Figure S3). The iron and acid labile sulfide content per mole of protein was 2.9 ± 0.1 and 4.6 ± 0.2, respectively. In an attempt to bolster the Fe/S content, chemical reconstitution with iron and sulfide was employed.^[24]^ Notably, increased amounts of iron and sulfur could be detected in the monomeric, reconstituted protein, 7.8 ± 0.2 and 10.8 ± 0.7, respectively. Consistent with increased iron/sulfur, the UV-vis spectrum showed an increase in the 320 and 420 nm signals (Figure S2B).

To characterize Nar1 in greater detail, Mössbauer spectroscopy was employed at various temperatures and magnetic fields using protein samples labeled with ^57^Fe. Nar1 was first prepared with ^57^Fe in the expression medium and analyzed as isolated under anaerobic conditions (^57^Fe labeling). The experimental data were simulated with two components.

Component 1 (Figure 2A, blue line, Table 1) had an isomer shift (*δ*) of 0.44 mms^-1^ and a quadrupole splitting (*ΔE_Q_*) of 1.22 mms^-1^. These parameters are characteristic of [4Fe-4S]^2+^ clusters coordinated by cysteine ligands.^[25–26]^ Unexpectedly, component 2 was simulated with *δ* = 0.26 mms^-1^ and *ΔE_Q_* = 0.60 mms^-1^, consistent with parameters for [2Fe-2S]^2+^ clusters (Figure 2A, red line and Table 1).^[27]^ Field-dependent Mössbauer spectroscopy at 4.2 K displayed magnetic hyperfine splitting, attributable solely to the external field, and unambiguously confirmed the presence of [2Fe-2S]^2+^ and [4Fe-4S]^2+^ clusters (Figure S4A-B and Table S1). To investigate which Fe/S clusters were introduced upon chemical reconstitution of the as-isolated protein, ^57^Fe was utilized in the chemical reconstitution reaction using as-isolated protein expressed in medium containing ^56^Fe (^56^Fe/^57^Fe labeling). Under these conditions, the experimental data could again be simulated with contributions from [4Fe-4S]^2+^ and [2Fe-2S]^2+^ clusters with slightly altered parameters as compared to the as-isolated protein (Figures 2B and S4C-D and Table 1). Simulation of the ^56^Fe/^57^Fe labeling data with parameters of the as-isolated [4Fe-4S]^2+^ cluster resulted in a slightly worse simulation (Figure S5A, Table S2), which suggested the presence of a [4Fe-4S]^2+^ cluster in an electronic environment somewhat distinct from that of the as-isolated sample. Small differences in isomer shift and quadruple splitting for the [2Fe-2S]^2+^ cluster suggested either the presence of the same [2Fe-2S] cluster as in the as-isolated sample, but in a slightly different electronic environment due to conformational changes caused by an additional [4Fe-4S] cluster, or a [2Fe-2S] cluster at a new binding site. In the former case, the presence of the same [2Fe-2S] cluster as in the as-isolated sample could result from the facile exchange of ^56^Fe for ^57^Fe at a solvent accessible [2Fe-2S] cluster.^[28]^ Lastly, the as-isolated sample prepared with ^57^Fe in the medium was also reconstituted in the presence of ^57^Fe so as to label all possible Fe/S clusters in Nar1 (^57^Fe/^57^Fe labelling, Figures 2C and S4E-F). The experimental data were simulated by two [4Fe-4S]^2+^ species with the same parameters as in the ^57^Fe and ^56^Fe/^57^Fe labeling schemes, in addition to a [2Fe-2S] species that closely matched that observed following ^56^Fe/Fe^57^ labeling (Figure 2C and Table 1). In addition to Fe/S clusters, another component was assigned as Fe^3+^ stemming from either unusual Fe isotope exchange of Fe/S clusters or the reconstitution reaction (Figures 2C and S4-5B). Therefore, Mössbauer spectroscopy traced the presence of at least two and possibly three distinct Fe/S clusters inserted either *in vivo* or *in vitro*.

**Table 1.**
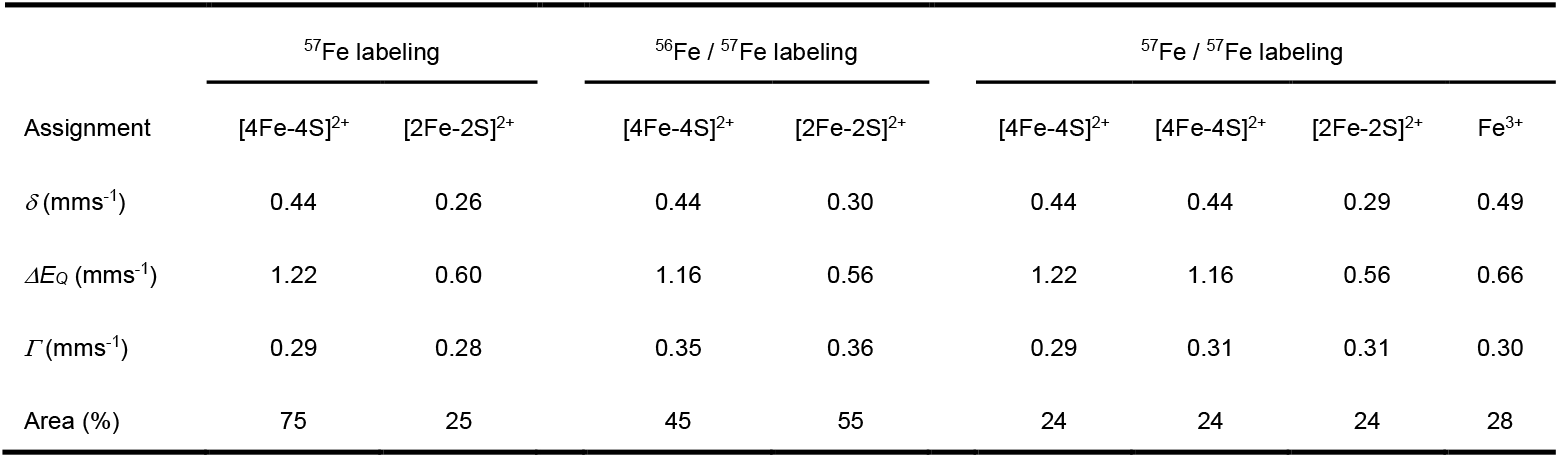
Parameters as obtained from the analysis of the Mössbauer spectra shown in Figure 2.

| Assignment | $^{57}\text{Fe}$ labeling | | $^{56}\text{Fe} / ^{57}\text{Fe}$ labeling | | $^{57}\text{Fe} / ^{57}\text{Fe}$ labeling | | | |
| --- | --- | --- | --- | --- | --- | --- | --- | --- |
| | [4Fe-4S]$^{2+}$ | [2Fe-2S]$^{2+}$ | [4Fe-4S]$^{2+}$ | [2Fe-2S]$^{2+}$ | [4Fe-4S]$^{2+}$ | [4Fe-4S]$^{2+}$ | [2Fe-2S]$^{2+}$ | $\text{Fe}^{3+}$ |
| $\delta$ (mm/s) | 0.44 | 0.26 | 0.44 | 0.30 | 0.44 | 0.44 | 0.29 | 0.49 |
| $\Delta E_Q$ (mm/s) | 1.22 | 0.60 | 1.16 | 0.56 | 1.22 | 1.16 | 0.56 | 0.66 |
| $\Gamma$ (mm/s) | 0.29 | 0.28 | 0.35 | 0.36 | 0.29 | 0.31 | 0.31 | 0.30 |
| Area (%) | 75 | 25 | 45 | 55 | 24 | 24 | 24 | 28 |

**Figure 2.**
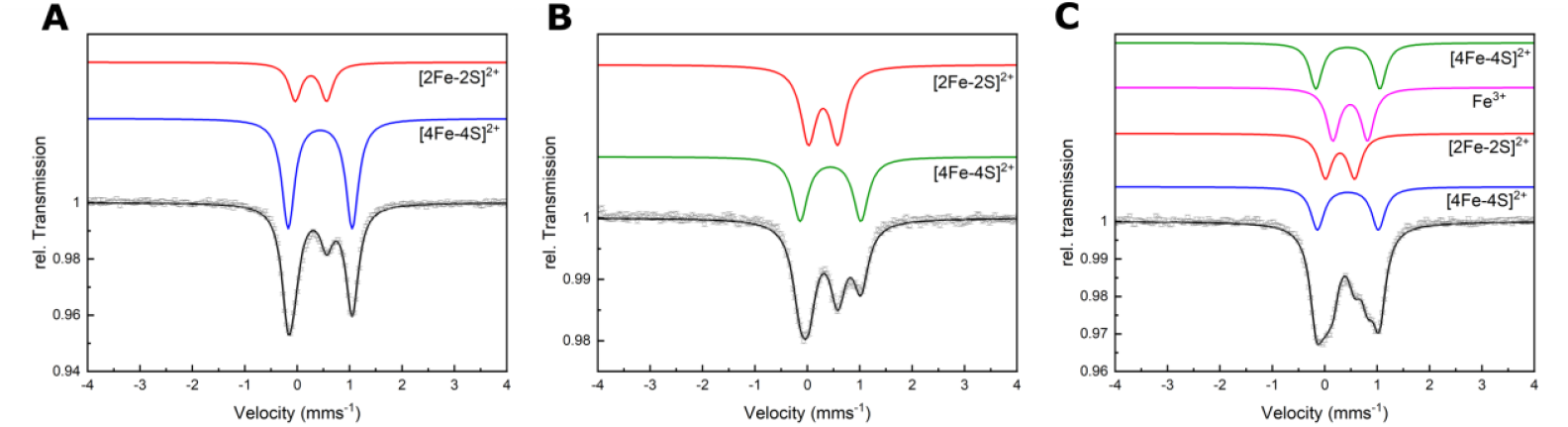
Mössbauer spectroscopy indicated the presence of [2Fe-2S] and [4Fe-4S] clusters bound to Nar1. A) Spectra of as-isolated Nar1 with ^57^Fe labeling, B) Nar1 with ^56^Fe/^57^Fe labeling, and C) Nar1 with ^57^Fe/^57^Fe labeling. Experimental data are shown as open circles with corresponding experimental error. Simulations are shown for the individually assigned components (in color) and of the experimental data combining the individual components (black). Spectra were collected at 77 K with protein concentrations ranging from 200-500 μM. For parameters see Table 1.

To gain more insight into the Fe/S clusters, we utilized variable-temperature EPR spectroscopy at 34 GHz/1.2 mT (Q-band). The Q-band spectrum of as-isolated Nar1 reduced with sodium dithionite (DT) showed a broad and complex signal at 40 – 60 K (Figure 3A, bottom trace and S6). This signal corresponds to slow relaxing low spin *S* = ½ [2Fe-2S]^+^ clusters^[29–30]^ and is more complicated than a single rhombic [2Fe-2S] cluster signature that is found in [2Fe-2S] ferredoxins.^[31–32]^ The Nar1 spectrum is reminiscent of multiple [2Fe-2S]^+^ components detected in BioB and Anamorsin,^[33–34]^, which suggested multiple [2Fe-2S] species (Figure 3A). Spectra acquired at lower temperatures down to 10 K showed progressive complexity and gain of some signals as the temperature was lowered. A rhombic-like signal at 10 K indicated the presence of one or more low spin *S* = ½ [4Fe-4S]^+^ clusters, which are not detectable above 20 K due to fast relaxation (Figure 3A).^[29–30]^ Notably, at 5 K the spectrum broadened significantly, indicative of magnetic exchange coupling between Fe/S clusters. Overall, EPR supports [4Fe-4S] and [2Fe-2S] clusters binding to as-isolated Nar1, consistent with the Mössbauer data.

**Figure 3.**
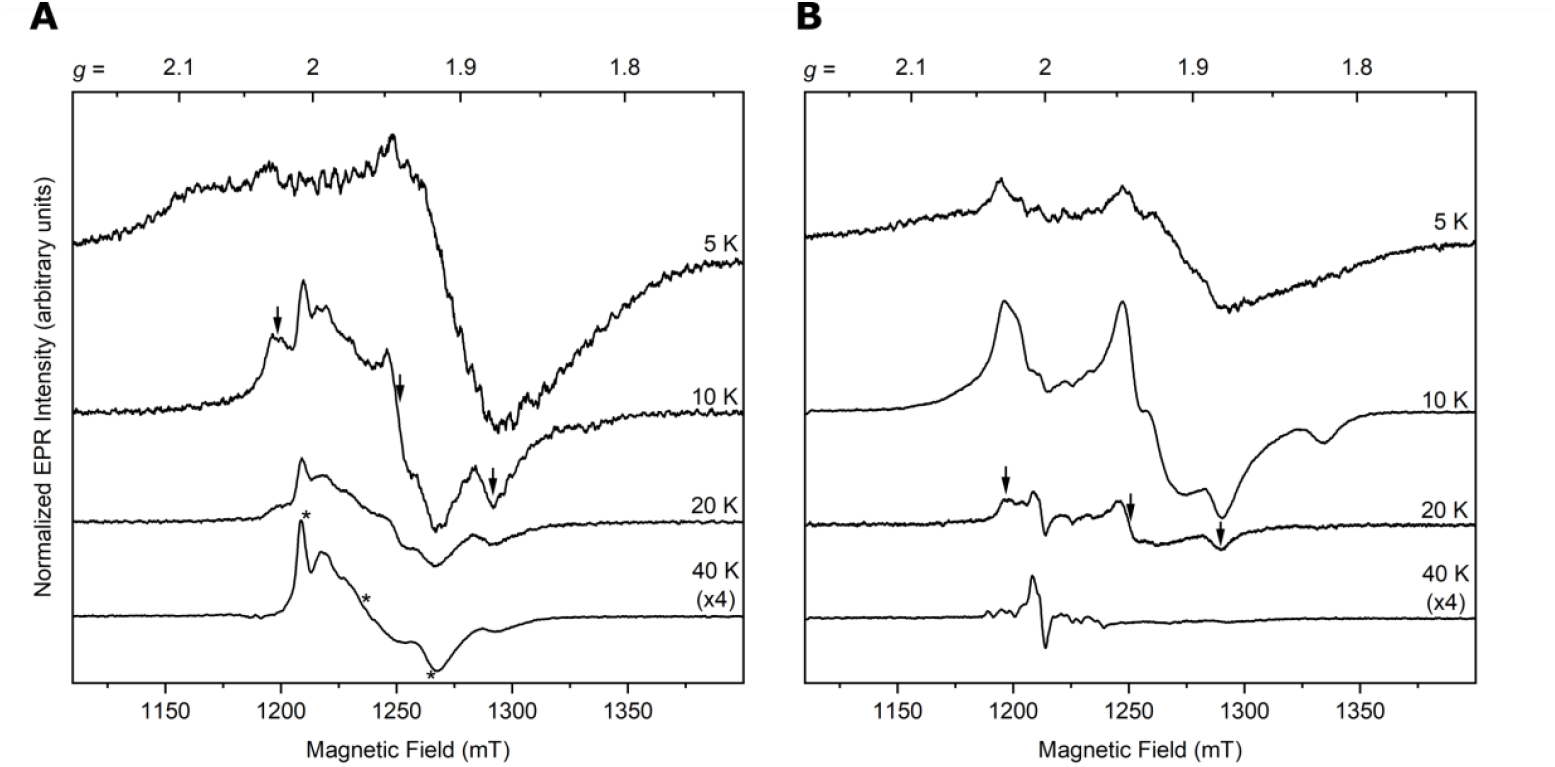
Temperature-dependent behavior of as-isolated (A, 930 μM) and reconstituted (B, 480 μM) Nar1 reduced with 10 equiv. of NaDT observed by pulsed Q-band EPR (34 GHz). The contributions of [2Fe-2S]^+^ clusters in as-isolated protein are seen at 40 K in (A), with a prominent signal comparable to *g* values for a [2Fe-2S]^+^ cluster in Anamorsin (marked by *).^[42]^ At 10 K in (A), a signal for a [4Fe-4S]^+^ cluster becomes apparent, which is comparable to the signal at 20 K in (B) for reconstituted protein (marked by arrows). The radical signal at 40 K in (B) likely stems from NaDT. All spectra are corrected for concentration and the 40 K spectra are multiplied by a factor of four.

Next, we investigated the reconstituted protein by EPR. The prominent [2Fe-2S]^+^ signals observed at higher temperatures in the as-isolated sample were absent (Figure 3B, bottom trace). Upon lowering the temperature to 20 K, a rhombic signature appeared with approximate *g* values of *g*1 ≈ 2.028, *g*2 ≈ 1.941, and *g*3 ≈ 1.882, similar to that observed with the as-isolated sample (Figure 3B). While these *g*-values are in the range of [4Fe-4S]^+^ clusters, they are different from axial and rhombic EPR signals observed previously for Nar1 and [FeFe]-hydrogenases.^[10–11, 16, 35–36]^ At 10 K, the spectrum broadened with additional features becoming apparent (*e*.*g*., at *g* ≈ 1.819) due most likely to at least one additional, faster-relaxing [4Fe-4S]^+^ cluster. The spectrum at 5 K was further broadened, again suggestive of magnetic coupling between [4Fe-4S]^+^ clusters (Figure 3B, top trace).^[35, 37–39]^ Although binuclear Fe/S clusters generally relax more slowly than tetranuclear Fe/S clusters, the presence of two [4Fe-4S] clusters in reconstituted Nar1 may promote fast relaxation of the [2Fe-2S] cluster^[40]^, making it difficult to detect at higher temperatures and to separate at low temperatures. Alternatively, the [2Fe-2S] cluster may have been destroyed by reduction with NaDT or converted into an EPR-inactive state, *e*.*g*., [2Fe-2S]^0^.^[41]^ We note that [2Fe-2S]^+^ and [4Fe-4S]^+^ clusters were not present in the non-reduced samples, only minor amounts of [3Fe-4S]^+^ clusters were apparent (Figure S7).

Finally, we used native mass spectrometry coupled with ion mobility (IMS-MS) to study the various species of Nar1 in the gas phase.^[43]^ Analyzing the folded species (Figure S3) showed Nar1 with various cofactor occupancies in the as-isolated protein. The highest abundance peak at 57,784 Da corresponded to Nar1 containing one [4Fe-4S] cluster (theoretical mass 57,785 Da), but protein bound to one [4Fe-4S] and one [2Fe-2S] cluster was also observed at a mass of 57,964 Da (Figure 4A, Table S3). Protein containing single [2Fe-2S] and [3Fe-4S] clusters were also observed in addition to a weak signal for apo protein. Measurement of the chemically reconstituted sample revealed additional peaks at 58,139 and 58,321 Da corresponding to [4Fe-4S]/[4Fe-4S]-Nar1 and [4Fe-4S]/[4Fe-4S]/[2Fe-2S]-Nar1, respectively (Figure 4B and Table S3). These observations not only verify a second [4Fe-4S] cluster binding to Nar1 but also a third site occupied by a [2Fe-2S] cluster, which was hinted at in the Mössbauer spectroscopy experiments. Additionally, disassembly of the [2Fe-2S] cluster leading to [4Fe-4S]/[Fe-S]-Nar1 and [4Fe-4S]/[4Fe-4S]/[Fe-S]-Nar1 adducts was observed by MS supporting the binding of mononuclear iron observed by Mössbauer spectroscopy (Figure S8 and Table S3). Both [4Fe-4S]/[4Fe-4S]-Nar1 and [4Fe-4S]/[4Fe-4S]/[2Fe-2S]-Nar1 could also be independently generated using a semi-biosynthetic reconstitution of Fe/S clusters employing the desulfurase NifS (Figure 4C).^[44]^ Strikingly, one of the [4Fe-4S] clusters in these states was highly prone to degradation by molecular oxygen under the experimental conditions (*t1/2* ≈ 9 min) whereas the singly occupied [4Fe-4S]-Nar1 species showed much greater stability (Figure 4C-D). The latter experiment indicated that the presence of Nar1 associated with only [2Fe-2S] or [3Fe-4S] clusters stemmed from the decay of [4Fe-4S] clusters (Figures 4 and S7-8).

**Figure 4.**
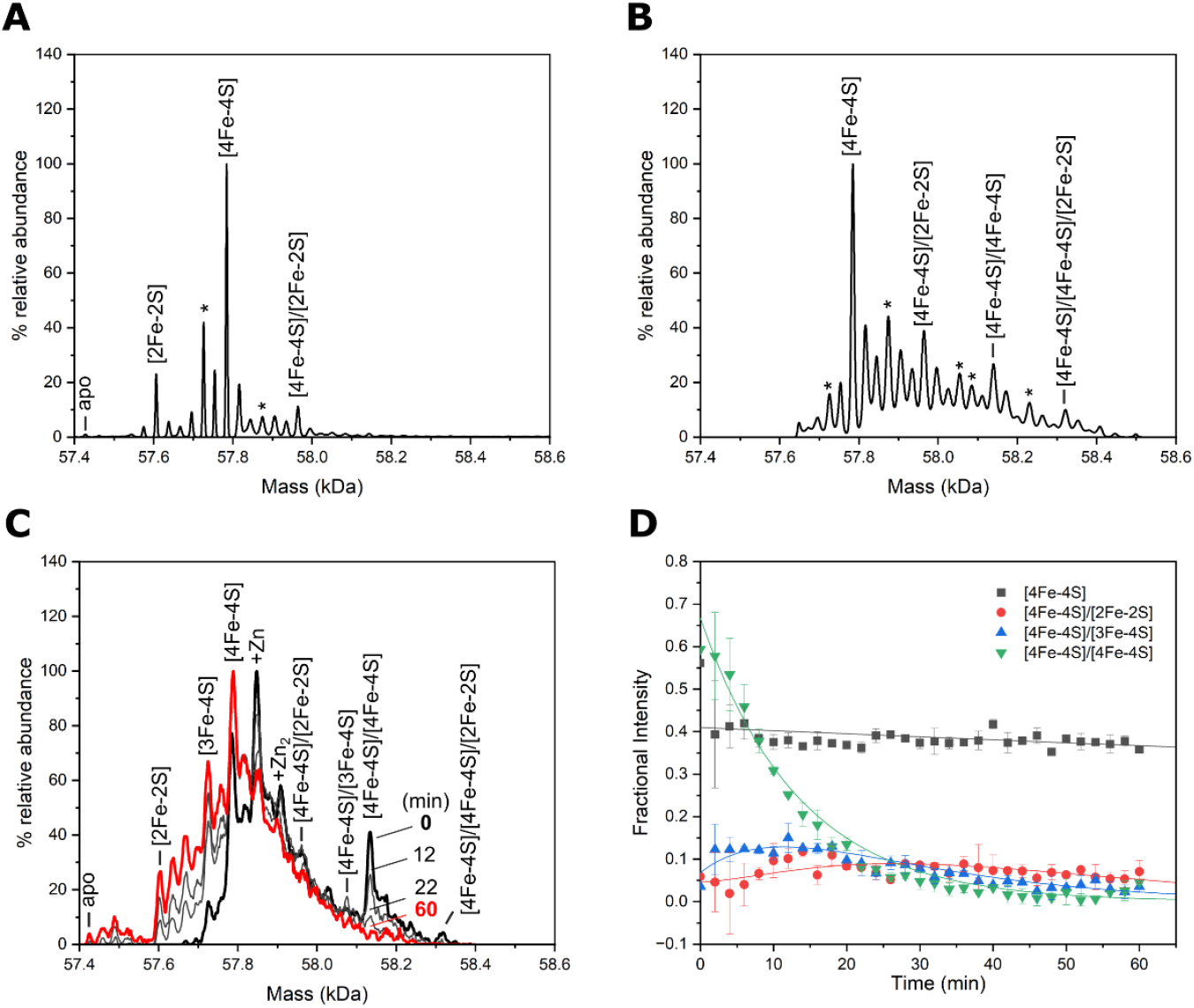
Native mass spectrometry of Nar1. A) As-isolated Nar1 binds up to two Fe/S clusters, with a [4Fe-4S] cluster predicted in site 1 and [2Fe-2S] cluster at an unknown site, in addition to singly occupied Fe/S species. B) Chemically reconstituted Nar1 binds an additional [4Fe-4S] cluster most likely in site 2, resulting in [4Fe-4S]/[4Fe-4S] and [4Fe-4S]/[4Fe-4S]/[2Fe-2S] species. Additional species are observed due to cluster breakdown (labeled with *) or sulfur adducts (see Figure S8). C) Spectra of enzymatically reconstituted Nar1 before (black line), and after exposure to dissolved atmospheric oxygen (60 min exposure; red line), measured on a lower resolution instrument set up for kinetics. Intervening spectra (grey lines), recorded at 12- and 22-min exposure show rapid loss of one of the [4Fe-4S] clusters. Cluster types are labelled in A-C based on molecular masses (Table S3) and analysis by spectroscopic methods. D) Temporal analysis of the [4Fe-4S] clusters in response to O_2_. One of the clusters in [4Fe-4S]/[4Fe-4S]-Nar1 rapidly degrades to unstable [3Fe-4S] and [2Fe-2S] clusters. We note that the [4Fe-4S]/[4Fe-4S]/[2Fe-2S]-Nar1 species under the conditions in (C) was too low in intensity to provide reliable kinetics. See Figure S9 for the temporal analysis of additional cluster degradation species.

Altogether, the biophysical data presented here provides new insight into the insertion of Fe/S clusters into the essential yeast protein Nar1 (Scheme 1). As the as-isolated protein contained both [2Fe-2S] and [4Fe-4S] clusters, it is proposed that recombinantly expressed Nar1 purifies from the *E. coli* Suf^++^ strain predominately with a [4Fe-4S] cluster at Fe/S site 1 as in *Chlamydomonas reinhardtii* HydA1 (Scheme 1)^[9, 11, 16, 36, 45]^ and at least one [2Fe-2S] cluster. The latter Fe/S cluster may have been critical for the chemical reconstitution of a second [4Fe-4S] cluster at Fe/S site 2. Iron and sulfide determinations indicated that in both as-isolated and reconstituted protein, not all sites are fully occupied, potentially due to either protein quality (conformations of apo protein incapable of cofactor binding) or lability of Fe/S clusters, as observed here and previously.^[9, 45]^ Yet, the spectroscopic and spectrometric data on reconstituted samples presented here clearly demonstrated the binding of a second [4Fe-4S] cluster in site 2 of Nar1.

Although the Suf^++^ strain facilitated the insertion of at least one [2Fe-2S] cluster into Nar1, our data, in addition to previous studies^[9–11, 16]^, suggest that the [4Fe-4S] cluster at Fe/S site 2 cannot be readily inserted *in vivo* by prokaryotic ISC or SUF machineries.^[46]^ The dedicated loading of the N-terminal [4Fe-4S] cluster *in vivo*^[9–10]^ therefore would be hypothesized to only be accomplished by the essential CIA scaffolding complexes involving Nbp35 or Cfd1/Nbp35 in eukaryotes (Scheme 1).^[7]^ This assumption would be consistent with the essentiality of Nbp35 in eukaryotes and its retention in the minimal CIA system of *Monocercomonoides exilis*.^[3, 47]^ *In vitro*, one of the [4Fe-4S] clusters, potentially also the [2Fe-2S] cluster, are highly susceptible to degradation by molecular oxygen. Intriguingly, genetic studies in various eukaryotes have also suggested a connection of Nar1 homologs to O2 sensitivity during cell growth.^[16, 48–50]^ Whether the Fe/S clusters of Nar1 has physiological roles in O2 sensing will be important questions to address in future *in vitro* and *in vivo* studies.

**Scheme 1.**
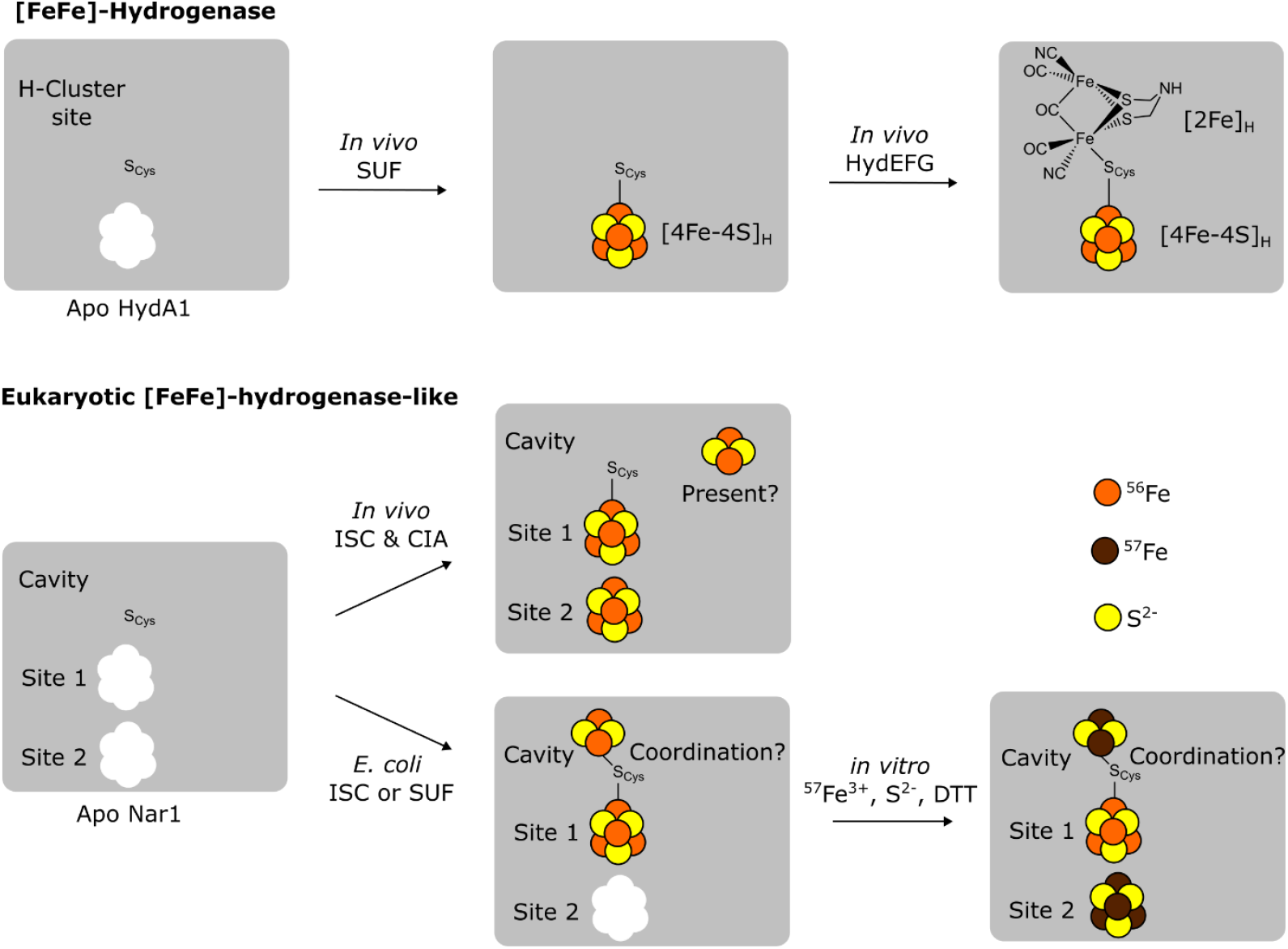
Models for the loading of cofactors into the homologous proteins [FeFe]-hydrogenase HydA1^[51]^ (top) and Nar1 (bottom). In [FeFe]-hydrogenases, a bridging cysteine (S_cys_) coordinates both the [4Fe-4S]_H_ cluster and the unique [2Fe]_H_ cluster. In addition to [4Fe-4S] clusters, Nar1 purified from *E. coil* is capable of binding at least one [2Fe-2S] cluster, which is hypothesized to be coordinated in an unknown fashion in the cavity of Nar1 that is analogous to the [2Fe]_H_ binding site.

Based on the Mössbauer and MS data together, a third binding site for a [2Fe-2S] cluster exists in Nar1. As there are no other obvious conserved Fe/S binding sites (Figure S1), it is hypothesized that the [2Fe-2S] cluster bound to Nar1 may occupy the cavity in which the [2Fe]H cluster resides in [FeFe]-hydrogenases, with an unknown coordination environment (Scheme 1). A [2Fe-2S] cluster binding at site 2 in as-isolated Nar1 may also be possible, yet such a cluster cannot explain the triply occupied species in reconstituted Nar1. The [2Fe-2S] cluster in Nar1 is most likely labile and/or solvent exposed as evidenced by exchange of Fe isotopes and its partial breakdown seen by MS. The complex EPR signature of the [2Fe-2S] cluster(s) in as-isolated protein, the relatively weak increase in UV-Vis absorptions signals in reconstituted protein, and the inability to detect the [2Fe-2S] cluster in reconstituted Nar1 by

EPR suggest a [2Fe-2S] cluster with unique properties. Further detailed spectroscopic and structure determination studies are underway to understand the binding of Fe/S clusters to Nar1. While it remains unclear if a [2Fe-2S] cluster is physiologically relevant in yeast and other eukaryotes, recent work showing the dependence of the [2Fe-2S] cluster-binding protein Apd1 on Nar1 and the CIA system warrants further studies.^[13, 52]^ Our study also now allows for the production of multiple Fe/S cluster-loaded forms of Nar1, which can be utilized in *in vitro* reconstitution assays to probe the molecular functions of the CIA pathway.

## Supporting Information

The authors have cited additional references within the Supporting Information.^[53-68]^

## Acknowledgements

J.J.B. and V.S. acknowledge financial support from the Deutsche Forschungsgemeinschaft (DFG)/German Research Foundation (SPP 1927, BR 5640/1-1, SCHU 1251/17-2). This work was also funded by the DFG under Germany’s Excellence Strategy (EXC 2033-390677874-RESOLV) to M.K. J.C.C. and N.L.B acknowledge funding from the UK’s Biotechnology and Biological Sciences Research Council (BB/V006851/1). We are grateful for contributions from Özge Efendi in protein preparation along with data collection and technical assistance from Marlina Indrayani, Steffi van anh Nguyen, and Martin Stümpfig. We thank Roland Lill for access to facilities for protein purification and acknowledge the contributions of the Core Facility ‘Protein Biochemistry and Spectroscopy’ of Philipps-Universität Marburg and Antonio J. Pierik for preliminary EPR experiments and discussion. We thank Sven Freibert for sharing the Nar1 plasmid and Patricia Kiley for the Suf^++^ *E. coli* strain. Molecular graphics were performed with UCSF Chimera, developed by the Resource for Biocomputing, Visualization, and Informatics at the University of California, San Francisco, with support from NIH P41-GM103311.

