## Supplemental Information for "Yeast [FeFe]-hydrogenase-like protein Nar1 can bind not only two [4Fe-4S] clusters but also a [2Fe-2S] cluster"

**Table of contents:**

1. **Supplemental Figures**

   **Figure S1**
   **Figure S2**
   **Figure S3**
   **Figure S4**
   **Figure S5**
   **Figure S6**
   **Figure S7**
   **Figure S8**
   **Figure S9**

2. **Supplemental Tables 1 - 3**

3. **Experimental Methods**

### Supplemental Figures

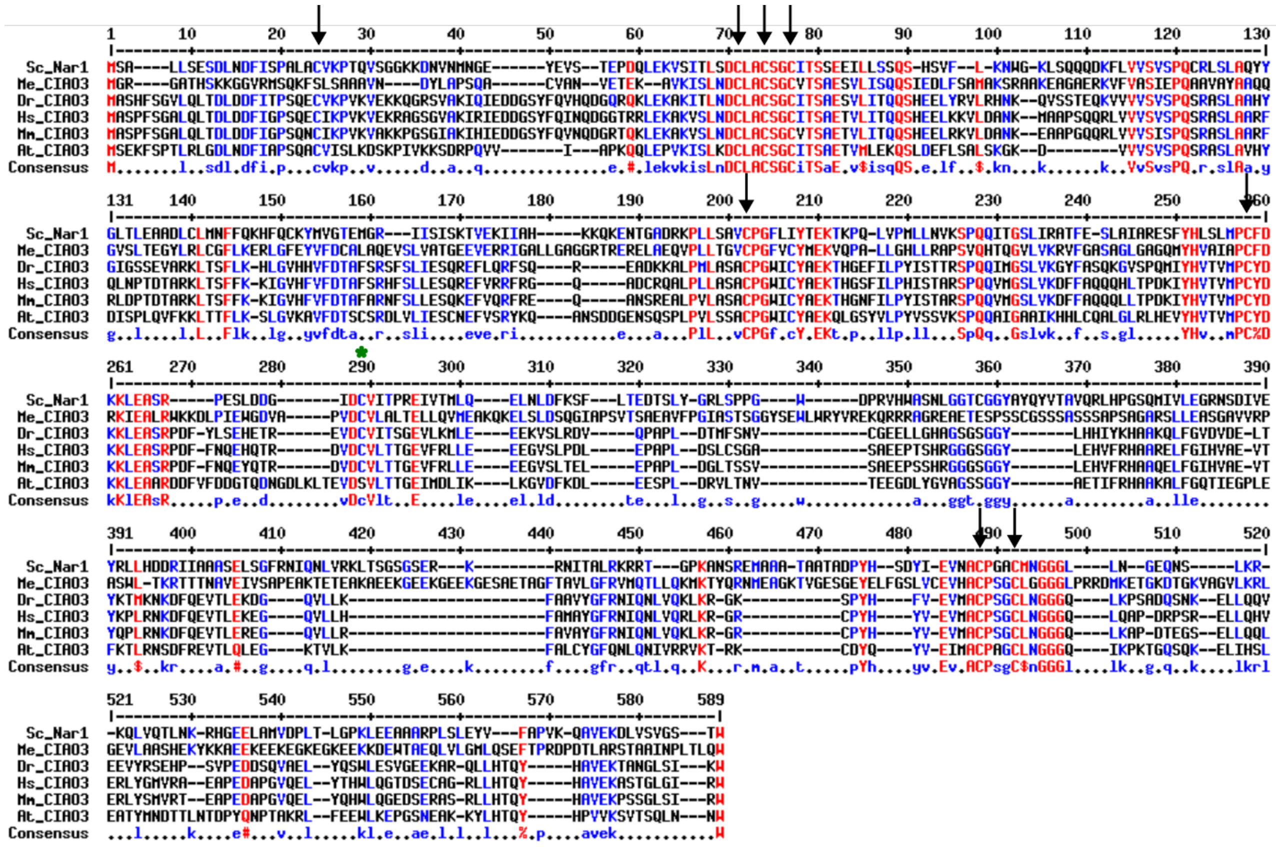

**Figure S1:** Multiple sequence alignment of Nar1 and select homologs from other eukaryotes. Arrows show the eight highly conserved cysteine residues involved in [4Fe-4S] cluster binding at sites 1 and 2 in Nar1. While one surface exposed cysteine is mostly conserved (green, *, from 1FEH model of Nar1, Figure 1); the remaining cysteines in Nar1 do not show any conservation. All homologs contain also a C-terminal tryptophan for targeting to the CTC. Sequences used were (organism, Uniprot or NCBI identifier): *Saccharomyces cerevisiae* (P23503), *Danio rerio* (A2RRV9), *Homo sapiens* (Q9H6Q4), *Mus musculus* (Q7TMW6), *Arabidopsis thaliana* (Q94CL6), *Monocercomonoides exilis* (XP_067724621). Alignment was made with the Dayhoff alignment parameters using MultAlin (http://multalin.toulouse.inra.fr/multalin/).[53]

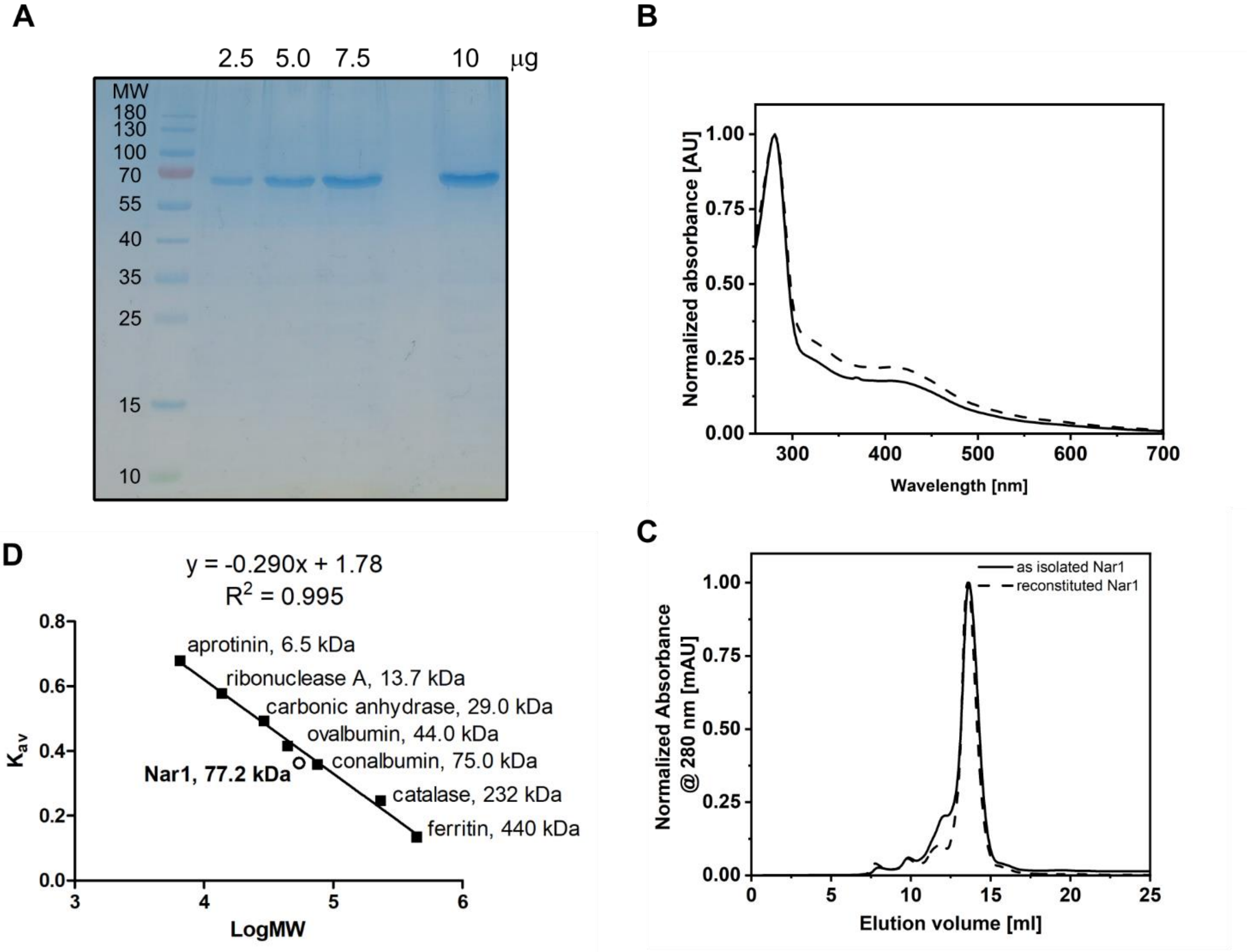


**Figure S2:** Purification of Nar1. A) SDS-PAGE of affinity-purified, as-isolated His-Nar1-Strep shown at multiple loading amounts. Molecular weight (MW, in kDa) ladder is shown on the left. B) As-isolated (solid line) and reconstituted Nar1 (dotted line) has UV-Vis absorption features typical of Fe/S clusters at 320 and 420 nm. Normalized UV-vis spectra in (B) at 280 nm correspond to the monomeric fraction of the SEC purified protein (C). C) HPLC-SEC analysis of as-isolated (solid line) and reconstituted (dotted line) Nar1 showing the predominant monomeric state of Nar1. D) Calibration curve for the SEC analysis and MW determination (Nar1, theoretical 57.6 kDa, observed 77.2 kDa). The linear regression is shown above the graph; the corresponding void volume ($V_o$) determined by elution of blue dextran was 7.7 mL. Column volume was 24 mL.

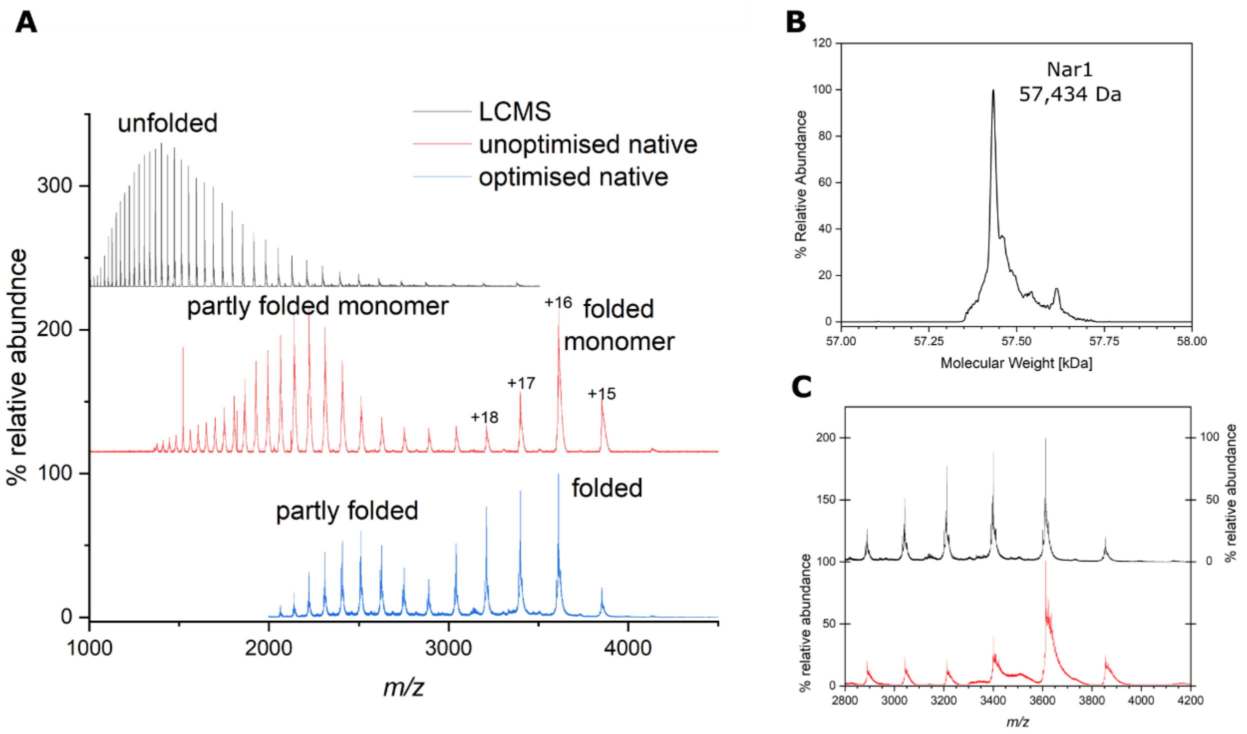


**Figure S3:** Mass spectrometric characterization of as-isolated Nar1. A) Positive mode ESI-TOF *m/z* spectra recorded under denaturing LCMS (black), and non-denaturing (native) MS conditions (red, blue). Minimizing in-source collision induced dissociation enhanced the transmission of the folded charge states for monomeric Nar1 (blue), relative to partially folded charge states (red) [54]. B) Deconvoluted mass spectrum of Nar1 corresponding to the *m/z* data in panel A. The theoretical molecular weight of His-Nar1-Strep is 57,567 Da and the observed molecular weight from LCMS data was 57,434 Da, indicating cleavage of the N-terminal Met by methionine aminopeptidase, giving a theoretical molecular weight 57,435 Da [55]. C) Comparison of the folded charge states for as-isolated (black), and reconstituted (red) Nar1 under native MS conditions.

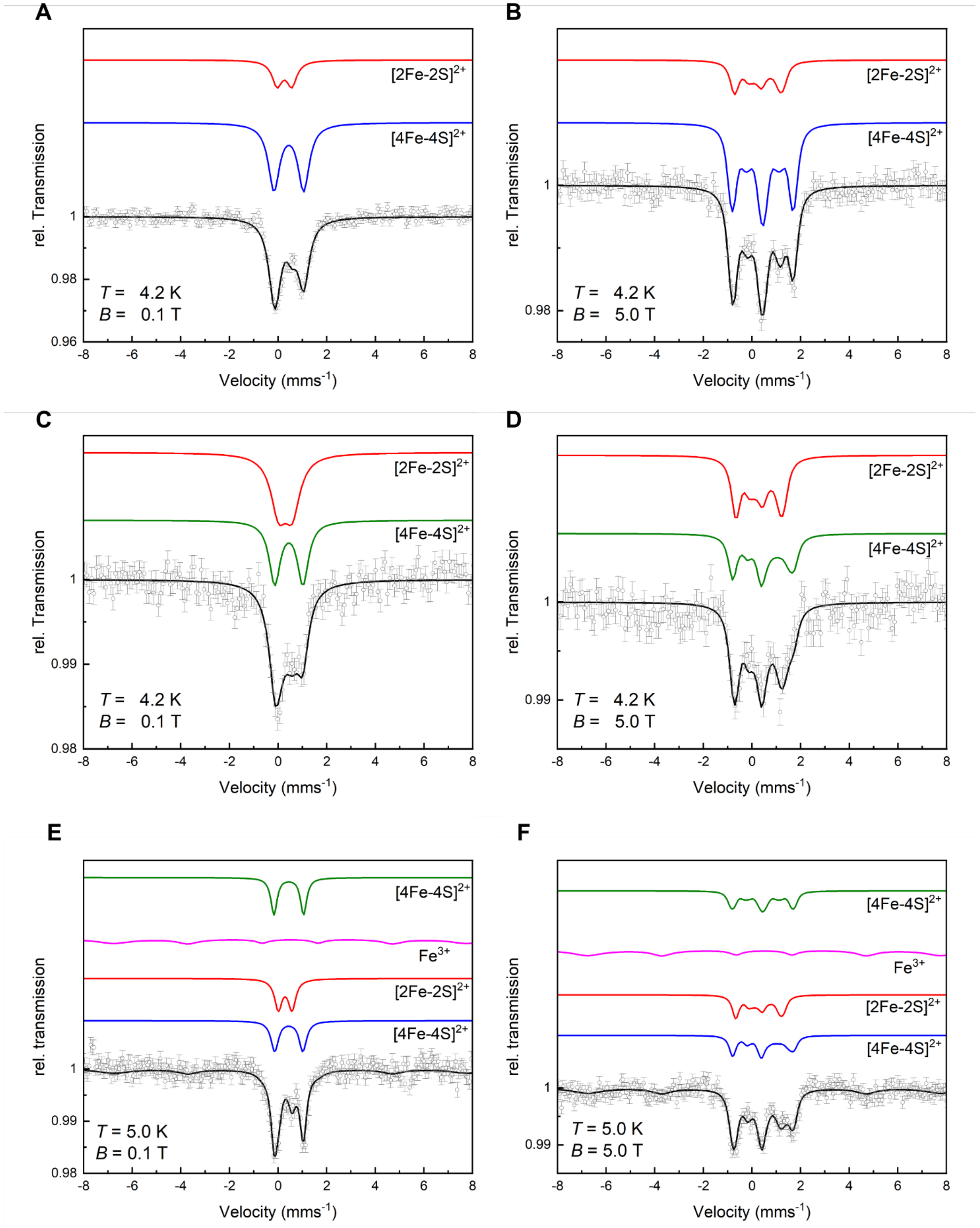


**Figure S4:** Applied field Mössbauer spectra on Nar1 in relation to Figure 3. The corresponding simulation parameters are listed in Table S1. The magnetic field was applied parallel to the γ-rays. Left panels show low-field data (0.1 T) and right panels show high-field data (5.0 T). Experimental data are shown as open circles with corresponding experimental error. Simulation of the individual components are shown in color with the sum of the corresponding components in black. A-B) Nar1 as-isolated using $^{57}Fe$ in the growth media, C-D) Nar1 as-isolated using $^{56}Fe$ in the growth media followed by reconstitution with $^{57}Fe$, and E-F) Nar1 as-isolated using $^{57}Fe$ in the growth media followed by reconstitution with $^{57}Fe$.

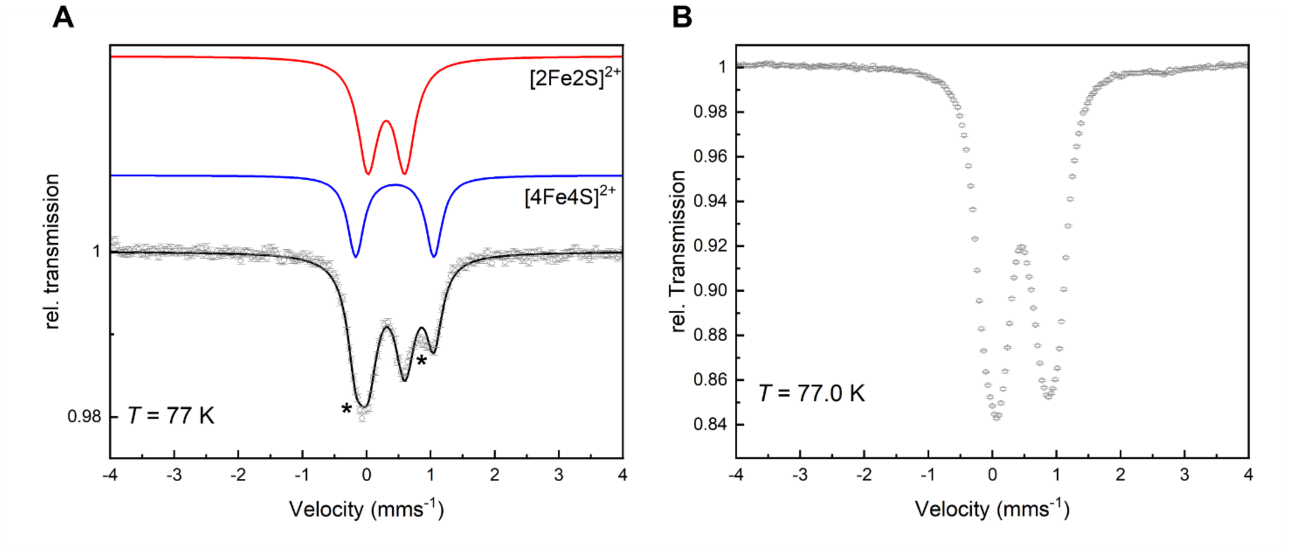


**Figure S5:** Supporting Mössbauer data for Figure 3. The corresponding simulation parameters are listed in Table S2. Data and simulations are represented as in Figure S4. A) Alternative simulation of as-isolated Nar1 using $^{56}Fe$ in the growth media followed by reconstitution with $^{57}Fe$ with the $[4Fe\text{-}4S]^{2+}$ components used for the simulation of the as-isolated protein with $^{57}Fe$ (compare with Figure 3B). * Asterisk denotes the deviation of the simulated fit as compared to the experimental data. The corresponding parameters are listed in Table S2. B) SEC purification of Nar1 is required after the chemical Fe/S cluster reconstitution assay. Mössbauer spectrum of as-isolated Nar1 followed by the reconstitution reaction with $^{57}Fe$ ($^{56}Fe/^{57}Fe$ labeling) and without further SEC purification. The Mössbauer spectrum is dominated by a signal with $\delta = 0.46$ mms$^{-1}$ (see Table S2) corresponding to non-specifically bound $Fe^{3+}$. SEC purification led to the removal of this species, as detected by Mössbauer spectroscopy (refer to Figure S4C-D).

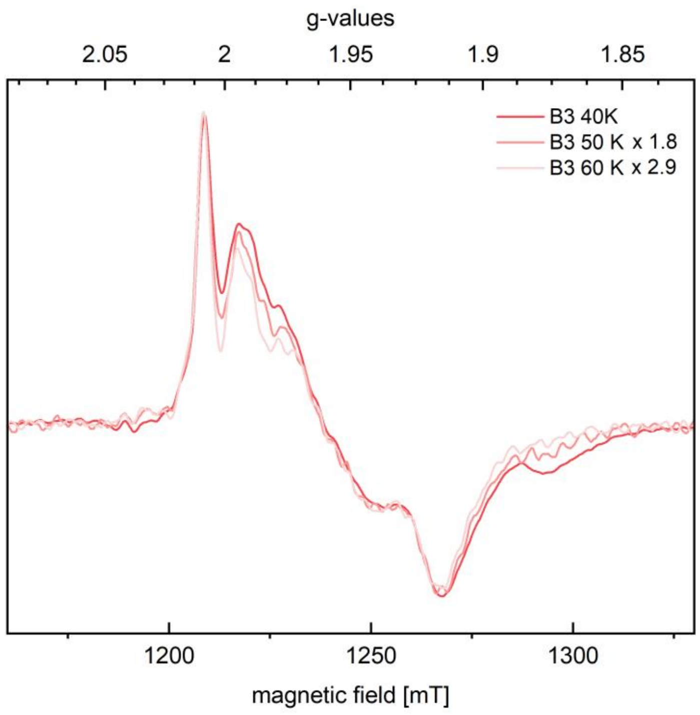


**Figure S6:** Pseudomodulated pulsed Q-band EPR spectra of as-isolated Nar1 at varying temperatures, in relation to Figure 4. Spectra are normalized to maximum. Temperatures are given in the inset.

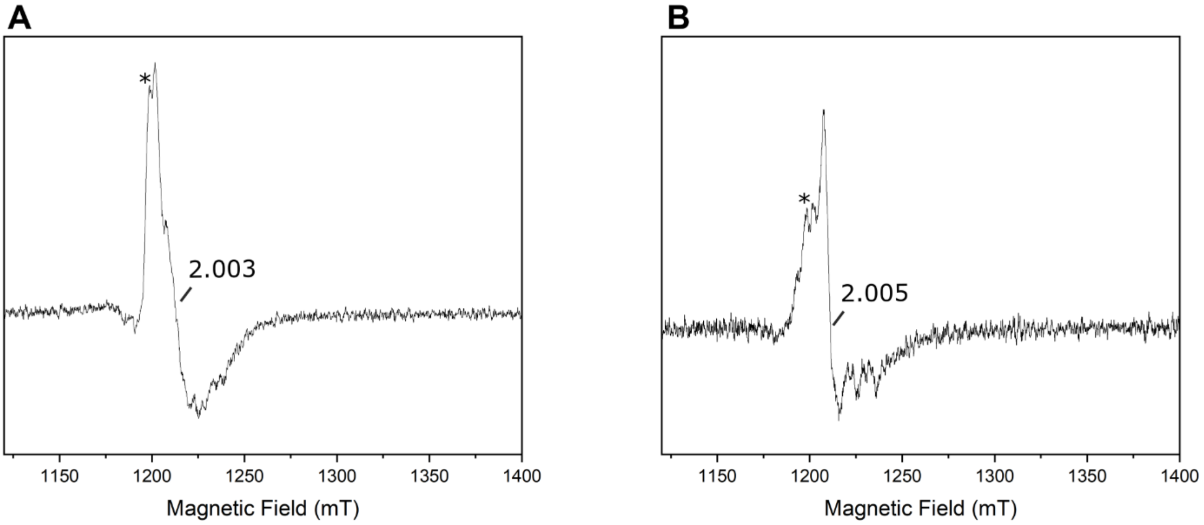


**Figure S7:** Pseudomodulated pulsed Q-band EPR spectra of non-reduced Nar1 samples at 10 K. In both as-isolated A) and reconstituted B) Nar1, minor amounts of a signal consistent with $[3Fe\text{-}4S]^{+}$ clusters were present at the indicated *g* values. . *Asterisk denotes the presence of $Mn^{2+}$ species.

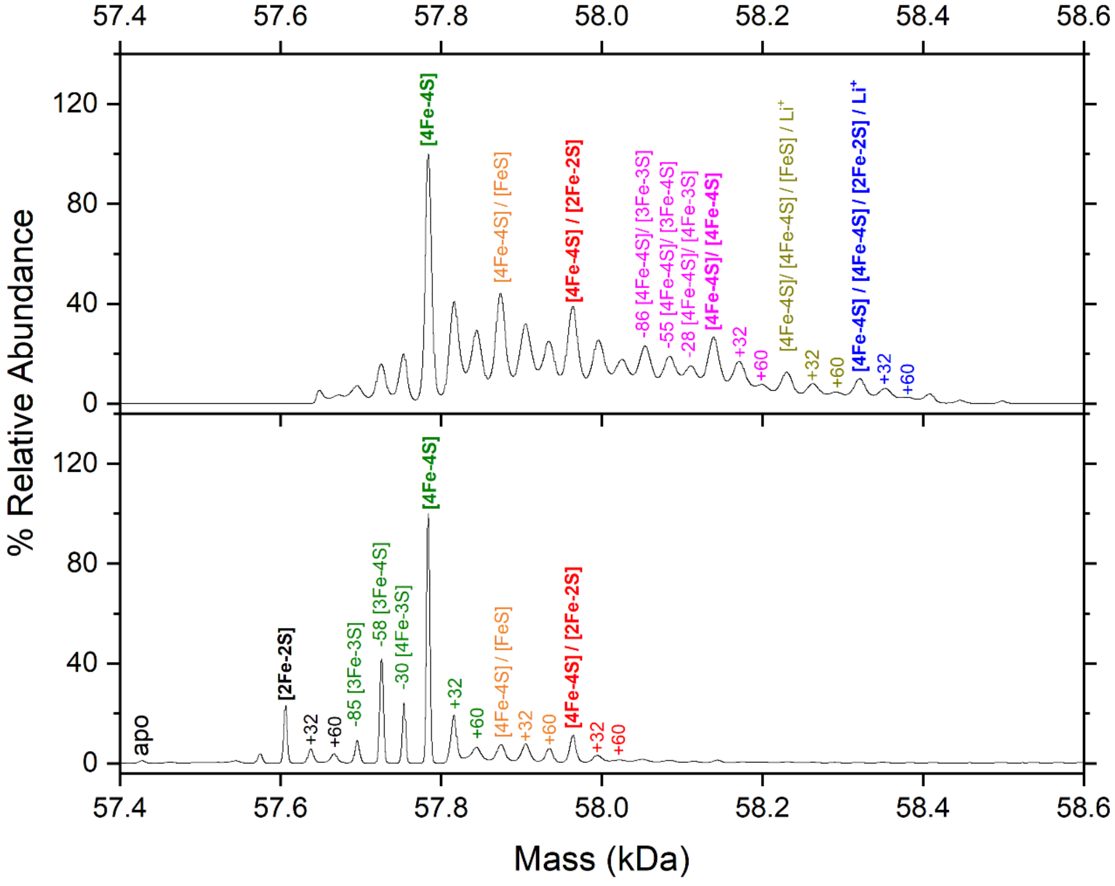


**Figure S8:** Assignment of additional peaks observed by native mass spectrometry for as-isolated (bottom) and reconstituted (top) Nar1 in reference to Figure 5A-B. Intact Fe/S clusters are labeled in bold (Table S3) and their corresponding decay or sulfur adducts are color coded based on species. Persulfide adducts (RS-$S^-$, +32 Da and RS-S-S-SR, +60 Da) were observed from as-isolated protein and in *in vitro* Fe/S reconstitution reactions. These sulfur adducts explain the elevated sulfide concentrations that were determined as compared to iron. Lithium ion adducts arise from the use of $Li_2S$ in reconstitution reactions.

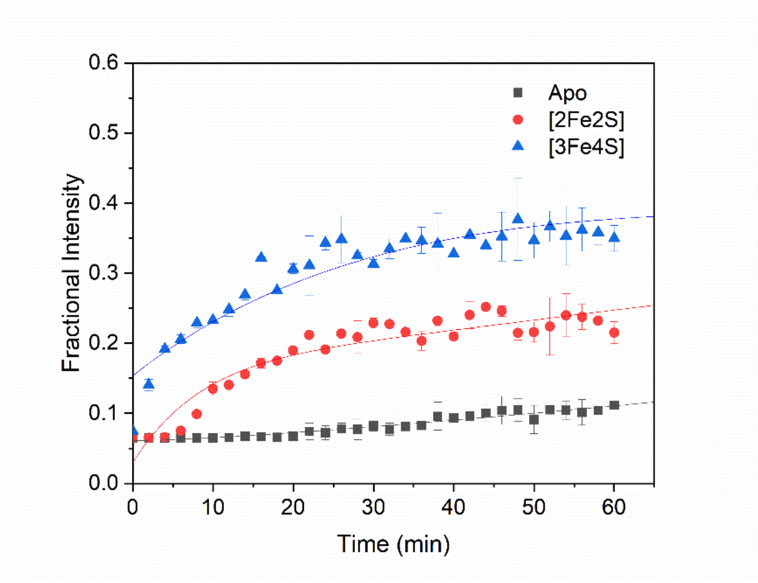


**Figure S9:** Temporal analysis of additional Nar1 species formed during the decomposition of Nar1 under aerobic conditions, in relation to Figure 5C-D. Apo Nar1 (squares) and singly bound [2Fe-2S] (circles) and [3Fe-4S] (triangles) cluster species increase in abundance as the [4Fe-4S] clusters in Sites 1 and 2 degrade.

### Supplemental Tables

**Table S1.** Mössbauer parameters of the simulations shown in Figure S4.

| | $^{57}Fe$ in Medium[a] | | $^{56}Fe$ in Medium / $^{57}Fe$ reconstitution[a] | | $^{57}Fe$ in Medium / $^{57}Fe$ reconstitution[b] | | | |
|---|---|---|---|---|---|---|---|---|
| **Assignment** | $[4Fe\text{-}4S]^{2+}$ | $2Fe\text{-}2S]^{2+}$ | $[4Fe\text{-}4S]^{2+}$ | $[2Fe\text{-}2S]^{2+}$ | $[4Fe\text{-}4S]^{2+}$ | $[4Fe\text{-}4S]^{2+}$ | $[2Fe\text{-}2S]^{2+}$ | $Fe^{3+}$[c] |
| **0.1 T** | | | | | | | | |
| $\delta$ (mms$^{-1}$) | 0.44 | 0.26 | 0.44 | 0.30 | 0.44 | 0.44 | 0.29 | 0.50 |
| $\Delta E_Q$ (mms$^{-1}$) | 1.22 | 0.60 | 1.16 | 0.56 | 1.16 | 1.22 | 0.56 | 0.00 |
| $\Gamma$ (mms$^{-1}$) | 0.55 | 0.50 | 0.55 | 0.77 | 0.37 | 0.30 | 0.37 | 0.6, 1.0 1.6 |
| Area (%) | 75 | 25 | 45 | 55 | 24 | 24 | 24 | 28 |
| **5.0 T** | | | | | | | | |
| $\delta$ (mms$^{-1}$) | 0.44 | 0.26 | 0.44 | 0.30 | 0.44 | 0.44 | 0.29 | 0.50 |
| $\Delta E_Q$ (mms$^{-1}$) | 1.22 | 0.60 | 1.16 | 0.56 | 1.16 | 1.22 | 0.56 | 0.00 |
| $\Gamma$ (mms$^{-1}$) | 0.36 | 0.40 | 0.40 | 0.41 | 0.31 | 0.30 | 0.31 | 0.6, 1.0, 1.5 |
| Area (%) | 75 | 25 | 45 | 55 | 50 | 24 | 24 | 28 |
| $\eta$[c] | 1 | 0 | 0 | 0 | 0 | 1 | 0 | 0 |

[a] Data collected at 4.2 K: [b] data collected at 5.0 K. [c] The $Fe^{3+}$ species stems from non-specific bound iron originating from the reconstitution reaction. The magnetic six-line pattern has been analyzed with a magnetic hyperfine field of 45 T. Refer to Figure S5B. [c] $\eta$, is the asymmetry parameter of the electric field gradient [56].

**Table S2:** Mössbauer parameters for the simulations shown in Figure S5.

| | $^{56}Fe$ in Medium / $^{57}Fe$ reconstitution Simulations for Figure S5A | | $^{56}Fe$ in Medium / $^{57}Fe$ reconstitution No SEC purification, Figure S5B |
|---|---|---|---|
| **Assignment** | $[4Fe\text{-}4S]^{2+}$ | $2Fe\text{-}2S]^{2+}$ | $Fe^{3+}$ |
| $\delta$ (mms$^{-1}$) | 0.44 | 0.31 | 0.46 |
| $\Delta E_Q$ (mms$^{-1}$) | 1.22 | 0.58 | 0.84 |
| $\Gamma$ (mms$^{-1}$) | 0.31 | 0.38 | 0.51 |
| Area (%) | 38 | 62 | 100 |

**Table S3:** Comparison of observed and theoretical masses for Nar1 species detected by native mass spectrometry.

| Species | Observed (Da) | Theoretical[a] (Da) | Species | Observed (Da) | Theoretical[a] (Da) |
|---|---|---|---|---|---|
| **As-isolated** | | | | | |
| apo | 57,427[b] | 57,435 | | | |
| [2Fe-2S]$^{2+}$ | 57,607 | 57,609 | | | |
| [3Fe-4S]$^{1+}$ | 57,726 | 57,730 | | | |
| [4Fe-4S]$^{2+}$ | 57,784 | 57,785 | | | |
| [4Fe-4S]$^{2+}$/[Fe(II)-S]<br>[4Fe-4S]$^{1+}$/[Fe(II)-S] | 57,875[c] | 57,874<br>57,875 | | | |
| [4Fe-4S]$^{2+}$/[2Fe-2S]$^{2+}$<br>[4Fe-4S]$^{2+}$/[2Fe-2S]$^{+}$<br>[4Fe-4S]$^{+}$/[2Fe-2S]$^{+}$<br>[4Fe-4S]$^{+}$/[2Fe-2S]$^{0}$ | 57,964[c] | 57,959<br>57,960<br>57,961<br>57,962 | | | |
| **Chemically reconstituted** | | | **Enzymatically reconstituted** | | |
| | | | apo | 57,427[a] | 57,435 |
| | | | [2Fe-2S]$^{2+}$ | 57,604 | 57,609 |
| [3Fe-4S]$^{1+}$ | 57,725 | 57,730 | [3Fe-4S]$^{1+}$ | 57,726 | 57,730 |
| [4Fe-4S]$^{2+}$ | 57,784 | 57,785 | [4Fe-4S]$^{2+}$ | 57,785 | 57,785 |
| [4Fe-4S]$^{2+}$/[Fe(II)-S]<br>[4Fe-4S]$^{1+}$/[Fe(II)-S] | 57,875[c] | 57,874<br>57,875 | [4Fe-4S]$^{2+}$ + $Zn^{2+}$ | 57,848 | 57,848 |
| | | | [4Fe-4S]$^{2+}$ + $(Zn^{2+})_2$[e] | 57,909 | 57,911 |
| [4Fe-4S]$^{2+}$/[2Fe-2S]$^{2+}$<br>[4Fe-4S]$^{2+}$/[2Fe-2S]$^{+}$<br>[4Fe-4S]$^{+}$/[2Fe-2S]$^{+}$<br>[4Fe-4S]$^{+}$/[2Fe-2S]$^{0}$ | 57,964[c] | 57,959<br>57,960<br>57,961<br>57,962 | [4Fe-4S]$^{2+}$/[2Fe-2S]$^{2+}$<br>[4Fe-4S]$^{2+}$/[2Fe-2S]$^{+}$<br>[4Fe-4S]$^{+}$/[2Fe-2S]$^{+}$<br>[4Fe-4S]$^{+}$/[2Fe-2S]$^{0}$ | 57,963[c] | 57,959<br>57,960<br>57,961<br>57,962 |
| [4Fe-4S]$^{2+}$/[3Fe-4S]$^{+}$<br>[4Fe-4S]$^{+}$/[3Fe-4S]$^{+}$<br>[4Fe-4S]$^{+}$/[3Fe-4S]$^{0}$ | 58,084[c] | 58,080<br>58,081<br>58,082 | [4Fe-4S]$^{2+}$/[3Fe-4S]$^{+}$<br>[4Fe-4S]$^{+}$/[3Fe-4S]$^{+}$<br>[4Fe-4S]$^{+}$/[3Fe-4S]$^{0}$ | 57,075[c] | 58,080<br>58,081<br>58,082 |
| [4Fe-4S]$^{2+}$/[4Fe-4S]$^{2+}$<br>[4Fe-4S]$^{2+}$/[4Fe-4S]$^{+}$<br>[4Fe-4S]$^{+}$/[4Fe-4S]$^{+}$ | 58,139[c] | 58,135<br>58,136<br>58,137 | [4Fe-4S]$^{2+}$/[4Fe-4S]$^{2+}$<br>[4Fe-4S]$^{2+}$/[4Fe-4S]$^{+}$<br>[4Fe-4S]$^{+}$/[4Fe-4S]$^{+}$ | 58,137[c] | 58,135<br>58,136<br>58,137 |
| [4Fe-4S]$^{2+}$/[4Fe-4S]$^{2+}$/[Fe(II)-S] + $Li^{+}$[d]<br>[4Fe-4S]$^{2+}$/[4Fe-4S]$^{+}$/[Fe(II)-S] + $Li^{+}$[d]<br>[4Fe-4S]$^{+}$/[4Fe-4S]$^{+}$/[Fe(II)-S] + $Li^{+}$[d] | 58,230[c] | 58,229<br>58,230<br>58,231 | | | |
| [4Fe-4S]$^{2+}$/[4Fe-4S]$^{2+}$/[2Fe-2S]$^{2+}$ + $Li^{+}$[d]<br>[4Fe-4S]$^{2+}$/[4Fe-4S]$^{2+}$/[2Fe-2S]$^{+}$ + $Li^{+}$[d]<br>[4Fe-4S]$^{2+}$/[4Fe-4S]$^{+}$/[2Fe-2S]$^{+}$ + $Li^{+}$[d]<br>[4Fe-4S]$^{+}$/[4Fe-4S]$^{+}$/[2Fe-2S]$^{+}$ + $Li^{+}$[d]<br>[4Fe-4S]$^{+}$/[4Fe-4S]$^{+}$/[2Fe-2S]$^{0}$ + $Li^{+}$[d] | 58,321[c] | 58,315<br>58,316<br>58,317<br>58,318<br>58,319 | [4Fe-4S]$^{2+}$/[4Fe-4S]$^{2+}$/[2Fe-2S]$^{2+}$<br>[4Fe-4S]$^{2+}$/[4Fe-4S]$^{2+}$/[2Fe-2S]$^{+}$<br>[4Fe-4S]$^{2+}$/[4Fe-4S]$^{+}$/[2Fe-2S]$^{+}$<br>[4Fe-4S]$^{+}$/[4Fe-4S]$^{+}$/[2Fe-2S]$^{+}$<br>[4Fe-4S]$^{+}$/[4Fe-4S]$^{+}$/[2Fe-2S]$^{0}$ | 58,317[c][f] | 58,309<br>58,310<br>58,311<br>58,312<br>58,313 |

[a] Molecular weight of His-Nar1-Strep with N-terminal methionine cleavage (Figure S3), [b] difference (- 8Da) may be explained by the presence of 4 disulfide bonds, [c] small differences (± 3Da) between observed and theoretical Nar1 masses may stem from protonation events associated with the cluster(s) [54, 57-58]. [d] presence of $Li^{+}$ stems from the chemical reconstitution reaction where $Li_2S$ was used. [e] $Zn^{2+}$ stems from the buffer and/or proteins used in the enzymatically reconstituted protein. In protein containing only one [4Fe-4S], an adduct with two bound Zn atoms may suggest that both the second [4Fe-4S] site and the [2Fe-2S] site are filled with Zn. [f] this is a tentative assignment as the signal is weak under these conditions and overlaps with other species.

### Experimental Methods

#### ***Recombinant Protein Expression and Purification***

Competent *E.coli* BL21(DE3)Suf$^{++}$ cells (gift from Patricia Kiley[59]) were transformed with a pASK-IBA43plus in which the *S. cerevisiae NAR1* gene was ligated to generate a plasmid encoding Nar1 with an N-terminal His-tag and a C-terminal strep-tag (Figure S3D, gift from Sven Freibert and Roland Lill). 100 mL preculture incubated for 4 h at 37 °C was used to inoculate a 2 L LB medium main culture, which was subsequently grown at 37 °C. At an $OD_{600}$ of 0.4, ferric ammonium citrate (FAC, Sigma-Aldrich) was added to a final concentration of 500 μM followed by induction of protein expression at an $OD_{600}$ of 0.6 by the addition of 150 μL of a 2 mg/mL anhydrotetracycline (Sigma Aldrich) solution in ethanol. In order to label Nar1 with $^{57}Fe$, the exact same expression procedure was carried out with $^{57}$Fe-ammonium citrate ($^{57}$FAC). Expression temperature was subsequently shifted to 18 °C and cultures incubated for 16 h overnight. Cells were harvested by 12 min centrifugation at 5,000 g. The following steps were carried out in an anaerobic chamber (Coy Labs). The cells were resuspended in anaerobic lysis buffer (50 mM Tris pH 8.5, 300 mM NaCl, 30 mM imidazole) to which 1 g of CelLytic™ Express (Sigma-Aldrich) per 10 g pellet was added. After 30 min the lysate was transferred to anaerobic centrifugation bottles and centrifuged at 16,000 g for 30 min. The supernatant was passed through a self-packed Ni-NTA (Thermo Fisher) column. Protein contaminants were removed by washing the column with 3 CV of Lysis buffer. The bound protein was eluted with elution buffer (50 mM Tris, pH 8.5, 300 mM NaCl, 250 mM imidazole). Immediately, the protein was passed through a PD-10 desalting column (Cytiva) to remove imidazole and rebuffered in storage buffer (50 mM Tris, pH 8.5, 150 mM NaCl, 5% (v/v) glycerol). The protein was further purified by size exclusion chromatography (SEC) using a HiLoad 16/600 200 μg column (Cytiva) equilibrated in storage buffer. Protein concentration was determined using the Bradford assay (Bio-rad). Protein aliquots in storage buffer were stored at – 80 °C in airtight vials.

#### ***Fe/S-cluster reconstitution***

*Chemical reconstitution* – Fe/S clusters were reconstituted chemically in an anaerobic chamber according to the published protocol of Freibert *et al.* [60]. Preliminary reduction of the protein in storage buffer was carried out with 6 equivalents of dithiothreitol (DTT) followed by 30 min incubation. A further 6 eq. of DTT were added and immediately 5 eq. of FAC (or $^{57}$FAC for Mössbauer samples) were titrated into the mixture. After 30 min of incubation, $Li_2S$ (Sigma-Aldrich) was added dropwise to the solution resulting in a dark brown solution. To remove excess iron and sulfur, the mixture was desalted (storage buffer) via a PD10 column and further purified via SEC on the HiLoad 16/600 200 pg column. The iron and sulfur content of Nar1 was determined colorimetrically by ferene - and acid-labile sulfide determination assays, as previously described [61]. In brief, at least three different concentrations of protein samples were analysed spectrophotometrically at 670 nm and 593 nm, and compared to calibration curves prepared, respectively with $Li_2S$ and $(NH_4)_2Fe(SO_4)_2$ as standards.

*NifS reconstitution* - Nar1 samples were *in vitro* reconstituted using NifS-catalysed reaction, as previously described [62]. Briefly, Nar1 was diluted to ~12 μM, treated with 1 mM DTT, 0.25 mM L-cysteine, 120 μM $(NH_4)_2Fe^{(II)}(SO_4)_2$, and ~0.3 μM NifS and incubated at an ambient temperature for ~2 h. The volume of the sample was increased 2-fold with buffer (50 mM Tris, 100 mM NaCl, 5% (v/v) glycerol, pH 8.0) and passed through a HiTrap SP column (Cytiva) to remove low molecular weight reactants and by-products. Bound Nar1 was eluted with 50 mM Tris, 2 M NaCl, 5% (v/v) glycerol, pH 8.0 [63].

***Spectroscopic and Mass Spectrometric Methods***

*HPLC - Size exclusion chromatography & UV-Vis spectroscopy (HPLC-SEC-UV-Vis)*

Nar1 was characterized via a HPLC-SEC-UV-vis setup using a calibrated and equilibrated Superdex 200 Increase 10/300 GL (Cytiva) analytical column attached to a DIONEX 3000 system (ThermoFisher) consisting of a DIONEX UltiMate 3000 UHPLC pump in line with a DIONEX UltiMate 3000 Diode Array Detector (Figures 2A and S1B-C). UV-Vis spectra spanning from 260 to 700 nm were collected at 2 Hz intervals with 1 s response time as the buffer exited the SEC column at a flow rate of 0.5 mL/min. Protein concentrations varied between 10 – 500 μM.

*EPR*

*Pulsed Q-band EPR* (electron spin echo detected EPR)- Protein samples with a concentration of 0.5-1 mM were reduced with 3 to 15 mM of NaDT in an anaerobic chamber and flash frozen with liquid $N_2$ after a constant incubation time of 3 min. The EPR spectra were recorded on a Bruker ELEXSYS E580 Q-band EPR spectrometer with an Oxford Instruments CF935 cryostat and Oxford Instruments MercuryiTC temperature controller and the Bruker ER 5106QT-2 resonator. For pulsed field-sweep experiments, a two-pulse Hahn spin echo sequence $\pi/2$–$\tau$–$\pi$–$\tau$–echo without phase cycling was used. The temperature was varied between 5 and 60 K. The $\pi/2$- and $\pi$-pulse lengths varied between 12–13 and 24–26 ns, while a constant interpulse delay $\tau$ of 210 ns was chosen. If not stated otherwise, the resulting absorption spectra were pseudomodulated with modulation amplitudes between 1 and 3 G, baseline-corrected and normalized to frequency, video gain, shots per point, number of scans and the respective modulation amplitude by using MATLAB (R2024a).

*Mössbauer spectroscopy*

The 77 K Mössbauer spectra were recorded with a conventional spectrometer from Wissel GmbH with $LN_2$ bath cryostat (Oxford Instruments) in transmission geometry. The isomer shifts are given relative to $\alpha$-Fe at room temperature. The spectra were analysed with the program Vinda [64]. The spectra were simulated by least-square fits using Lorentzian line shapes. The low and high field spectra were recorded with a closed-cycle cryostat equipped with a superconducting magnet (CRYO Industries of America Inc.) with the applied field parallel to the $\gamma$-rays. The magnetically split spectra were simulated with the spin Hamiltonian formalism [65].

*Mass spectrometry*

*ESI-MS under non-denaturing conditions* - Samples of Nar1 (as-isolated or reconstituted) were exchanged into 100 mM ammonium formate pH 8.5 using PD Minitrap G-25 columns (Cytiva). Samples (~10 $\mu$M) were transferred from the anaerobic cabinet using gas tight syringes and infused directly into the ESI-source of the mass spectrometer operating in positive mode with a capillary voltage of 3.5 kV. The $O_2$ sensitivity of NifS-reconstituted Nar1 was determined by combining aliquots of aerobic and anaerobic ammonium acetate, just prior to infusion, as previously described [66]. Two different mass spectrometers were used: a Bruker micrOTOF-QIII (Bruker Daltonics) or a Water Synapt XS [Date reported in Figure 5A-B were recorded on the Water Synapt XS operating in IMS mode; Data reported in Figure 4C were recorded on the Bruker micrOTOF-QIII]. Parameters for the transmission of Nar1 were optimized according to Lagnowsky *et al* [67] or Crack *et al* [54] for each spectrometer. The instruments were calibrated with ESI-L low concentration tuning mix (Agilent Tech.) and/or sodium iodide (Waters Corp.). Data were acquired over the 1,000 – 6,000 m/z range for 5 min.

Processing and analysis of MS experimental data was carried out using Bruker compass data analysis v4.1 (Bruker Daltonik) or Waters Mass Lynx 4.2 with Drift-scope v2.9 (Waters Corp.). For IMS data, charge states belonging to the 'folded' region were selected and exported to Mass Lynx for processing. Neutral mass spectra were generated from *m/z* spectra using the maximum entropy deconvolution algorithm of each analysis suit, between 50 and 65 kDa. For $O_2$ sensitivity of Nar1, the fractional intensities were calculated from deconvoluted spectra, and the temporal data fitted using Dynafit (Biokin) [66, 68].
